# Metabolic processes shape microbial interaction distributions

**DOI:** 10.64898/2026.07.15.738716

**Authors:** Prajwal Padmanabha, Sara Mitri

## Abstract

Microbial communities are largely driven by metabolic interactions, whereby species compete for shared resources and exchange byproducts. Such interactions are commonly classified by their sign alone. Yet, theory shows that the strength of interactions and the shape of their distribution impact how many and which species coexist. What this shape looks like in microbial communities and why has rarely been examined. Starting from a consumer-resource model of resource competition and cross-feeding, we find that these metabolic processes generate a skewed distribution with “many weak, few strong” interactions – a pattern previously documented in the food webs of larger organisms. This skew emerges whenever species have different substrate preferences and when metabolite leakage is sufficiently high. Across nine microbial datasets spanning diverse community origins, empirical interactions not only qualitatively display this skewed pattern but quantitatively follow the relationship between higher-order statistics predicted by the model. Generalized Lotka–Volterra (gLV) models, the standard framework for predicting community diversity, typically assume Gaussian interaction strengths. Instead, we show that the observed “many weak, few strong” interaction distribution is better captured by a lognormal distribution than the symmetric Gaussian distribution. Sampling interactions in gLV models from a lognormal distribution yields more stable communities in simulations and more accurately predicts the diversity observed in the two datasets where community-assembly experiments were performed. Together, these results reveal a metabolic origin for the “many weak, few strong” pattern of microbial interactions and show that using empirically grounded interaction distributions improves predictions of community diversity.

**Significance:** How strongly do microbes affect each other’s growth? The distribution of their interaction strengths shapes how many species coexist in a community and how stable they are. Across nine datasets, we show that most interactions are weak, with rare strong outliers, mirroring the macro-organism “many weak, few strong” interaction pattern. Using mathematical models, we show that competition for shared resources and metabolite cross-feeding, both prevalent in microbes, suffice to generate this pattern. Accounting for the full distribution of interaction strengths, rather than only their mean and variance, improves predictions of community diversity in simulations and empirical datasets. This work connects microbial metabolic processes to community-level structure and provides a route for predicting diversity when interactions cannot be exhaustively measured.

## I. INTRODUCTION

The diversity and stability of microbial communities is key to their functioning. Host-associated microbiomes are a great example of how a diverse and stable ecosystem provides benefits to the host. But disturbances to this stable state can lead to dysbiosis [1]. Similarly, in industrial bioreactors, achieving and maintaining stable communities is necessary to ensure functional robustness [2]. Across many such examples, it is important to understand what makes a community stable. When will it maintain its diversity or be dominated by a few species? The answer depends on the interactions in the community – the effects one species has on the growth and death of another. While much attention has been paid to the sign of interactions (negative, positive, or neutral) that dominates in microbial communities [3–6], far less has been placed on how strong they are and how those strengths are distributed. Ecological theory has shown that both the magnitude and the distribution of interaction strengths strongly shape community diversity and stability [7]: Stronger or more variable interactions constrain how many species can stably coexist [8, 9], while the distribution of those strengths determines patterns of abundance and persistence [10–12]. Despite this, we know little about these properties in empirical microbial communities or how they arise.

One domain where the shape of interaction distributions is well characterized is that of macro-organism food webs. Here, interaction strengths follow a consistent pattern: most interactions are weak, and only a few are strong. This “many weak, few strong” pattern has been linked to stability and persistence properties of diverse communities [12, 13]. In these well studied systems, the pattern is attributed to body-size allometry, where interaction strength scales with mass ratios [14], or emerges when energetic and feasibility constraints are imposed on trophic predator-prey models [15]. However, in microbes, interactions are predominantly mediated by metabolite exchange [16] and the expected macrobial mechanisms might not obviously apply. Here we ask whether the same pattern arises in microbial systems. If so, what generates it? And what are its consequences for community stability and diversity?

Answering these questions requires linking the metabolic processes underlying microbial interactions to the distribution of interaction strengths they produce, and in turn linking that distribution to consequences for community diversity and stability. Consumer-resource models provide a natural starting point, where instead of specifying interactions directly, they describe how species consume, produce, and compete for chemical resources in their environment [17]. These models capture the resource competition and cross-feeding that are prevalent in microbial communities [18– 20]. How the distribution of interaction strengths shapes community diversity and stability has been extensively studied using generalized Lotka Volterra (gLV) models [16, 21]. However, interactions in these models lack a mechanistic basis [22], are fixed across environments, and are usually assumed to be Gaussian distributed, largely for mathematical convenience [8, 23]. Mapping consumer-resource models onto their equivalent gLV interactions [24] provides a mechanistic grounding for the interactions and their distribution, while retaining the power of the gLV framework to study community diversity and stability.

Here, we start from consumer-resource models that consider two processes: resource competition, which sets limits for coexistence based on resource diversity [17], and cross-feeding, which typically increases coexistence [18]. When we translate these models into gLV interactions using a previously published method [25], we observe predominantly negative values, a strong skew toward zero, and more outliers than expected, mirroring the “many weak, few strong” pattern observed in food webs of larger organisms [26]. Across nine empirical microbial datasets spanning diverse community origins and inference methods, measured interactions show the same signature of being consistently weak and predominantly negative. Beyond this qualitative match, their higher-order statistics (skewness and kurtosis) follow the quantitative scaling relationship our model predicts. We analytically show how resource competition and cross-feeding contribute to the skew, providing a metabolic explanation for the observed structure that better resembles a lognormal than Gaussian distribution. Finally, we show that parameterizing gLV models with lognormally distributed interactions derived from consumer-resource communities reduces the unbounded growth that commonly afflicts Gaussian-parameterized models and more accurately predicts community diversity in two of the nine datasets where community assembly experiments were performed [27, 28].

## II. RESULTS

### Resource competition and cross-feeding shape the distribution of interactions

We begin with a consumer-resource model where we consider two kinds of metabolic processes: competition for shared resources and cross-feeding through metabolite leakage. The total resource taken up is divided into usage for increasing biomass and metabolite leakage. In a chemostat environment with dilution rate *δ*, the dynamics of the abundance of species *i* (*n*_*i*_) and the concentration of resource *α* (*c* _*α*_) are:

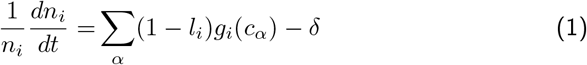

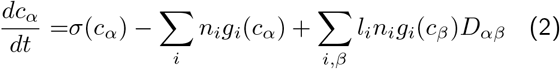

where for species *i, l*_*i*_ is the leakage fraction which represents the resources not used for growth, and the per-capita growth rate *g*_*i*_(*c*_*α*_) = *R*_*iα*_*c*_*α*_*/*(*K* + *c*_*α*_) (*R*_*iα*_ is the maximal growth rate and *K* is the Monod constant); for resource *α, σ*(*c*_*α*_) is the resource supply, and *D*_*αβ*_ is the stoichiometric resource conversion matrix (see Methods), similar to cross-feeding matrices in previous models [29]. We simulate the consumer-resource model until stationarity and use the final abundances of resources to calculate an effective growth rate and interaction coefficients for all species using previously published Environment-Organism (EO) framework ([25], see Methods and Table I for definition of the terms, Fig 1A). This method is rigorously equivalent to an accelerational framework [30] but the matrices it produces can be rescaled to obtain an effective gLV model at stationarity (see Methods). We note that there are many alternative approaches to obtain effective interaction matrices [17, 31, 32], which we discuss in supplement S1.

**TABLE I.**
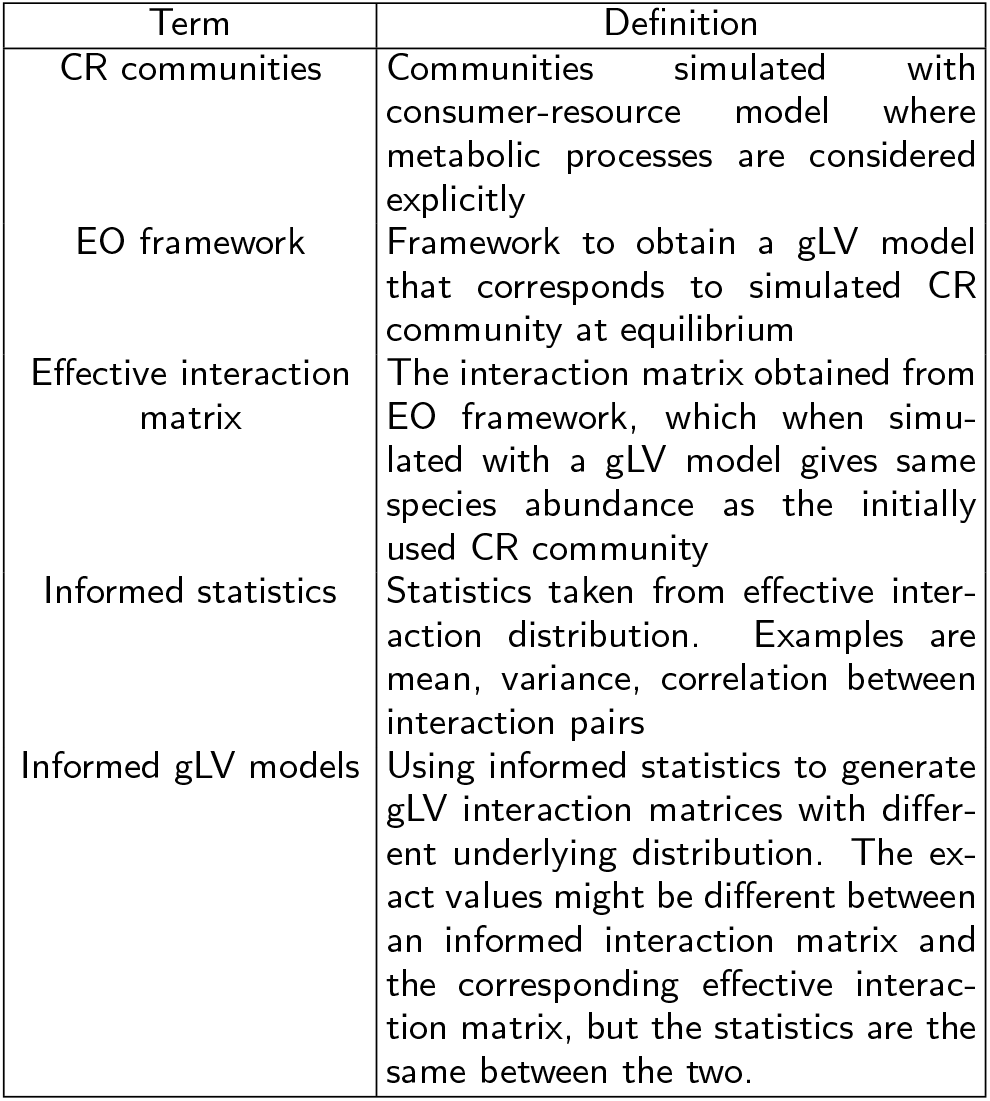
Definitions of terms used in the paper.

| Term | Definition |
| --- | --- |
| CR communities | Communities simulated with consumer-resource model where metabolic processes are considered explicitly |
| EO framework | Framework to obtain a gLV model that corresponds to simulated CR community at equilibrium |
| Effective interaction matrix | The interaction matrix obtained from EO framework, which when simulated with a gLV model gives same species abundance as the initially used CR community |
| Informed statistics | Statistics taken from effective interaction distribution. Examples are mean, variance, correlation between interaction pairs |
| Informed gLV models | Using informed statistics to generate gLV interaction matrices with different underlying distribution. The exact values might be different between an informed interaction matrix and the corresponding effective interaction matrix, but the statistics are the same between the two. |

**FIG. 1.**
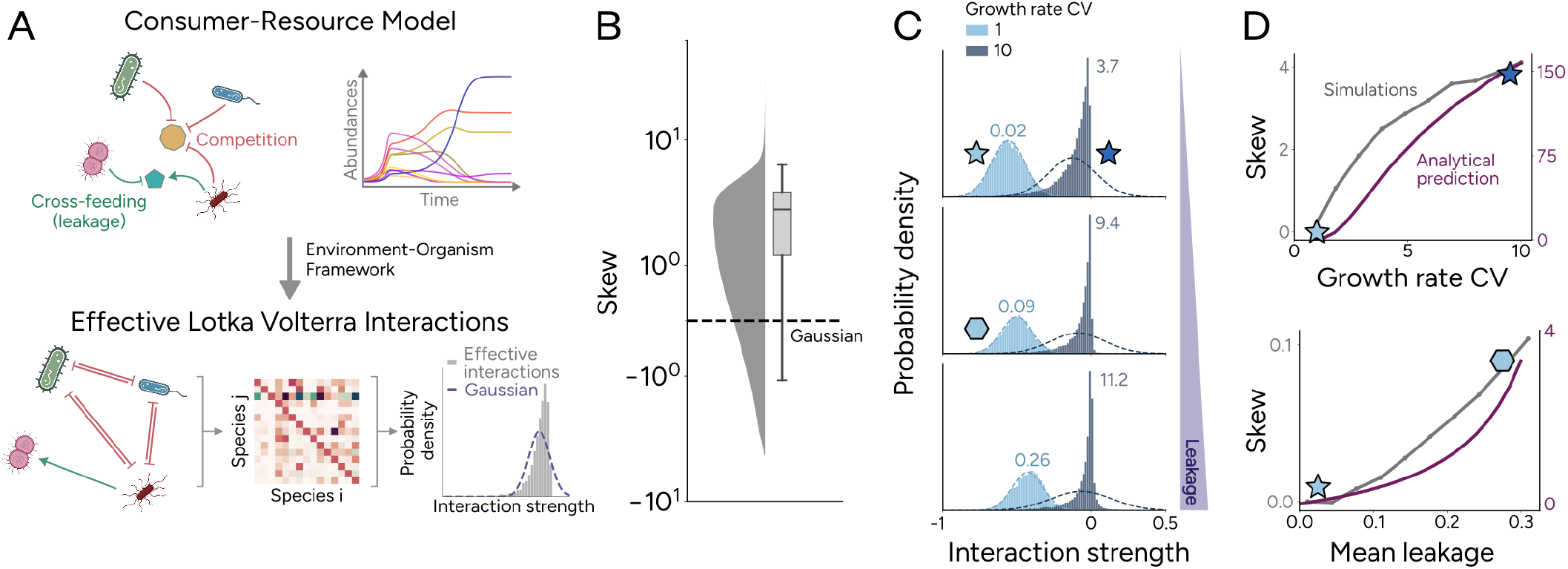
Effective interactions from consumer-resource communities. **A)** Schematic showing how resource competition and cross-feeding generate effective gLV interactions from consumer-resource models. Species abundances are simulated to steady state, at which point effective gLV interactions exactly capture the equilibrium abundances. An example distribution of effective interaction values (gray) is shown alongside a Gaussian with matched mean and variance (dashed) **B)** Violin and box plots showing skew values from 1000 simulated consumer-resource communities across various community sizes and metabolic parameters. Dashed line indicates the Gaussian expectation of zero. **C)** Example interaction distributions at increasing mean leakage levels (0.025, 0.275, and 0.575 from top to bottom). Within each panel, different colored histograms correspond to different growth rate coefficients of variation (CV, or ratio of standard deviation to mean; light blue = 1, dark blue = 10). Numbers above distributions indicate their skew. **D)** Distribution skew increasing as a function of: (top) growth rate heterogeneity, measured as the CV of *R*_*iα*_, and (bottom) mean leakage fraction. Gray lines show simulation results and purple lines show analytical predictions from a simplified model (see S5). Analytical predictions, which assume small leakage rates, capture qualitative trends rather than exact values. Colored symbols represent the distributions in panel C.

Using our consumer-resource model, we simulated 1000 communities of various community sizes, each assembled to stationarity with species having independently drawn metabolic parameters (see Methods). To quantify the deviation from normality, we measured the skewness of the interaction distributions, defined as the ratio of third cumulant to the cube of the standard deviation (see Methods). This metric has been widely used in food web studies to characterize the asymmetry of interaction strength distributions [13, 26]. Larger positive values indicate that interactions cluster near zero, with a long tail of stronger values, which gives the classic “many weak, few strong” patterns. The 1000 simulated communities have skewness values significantly different from the Gaussian expectation of zero (Wilcoxon signed rank *p <* 0.001, Figure 1B). Depending on the metabolic parameters chosen, the degree of skewness varies from nearly Gaussian at low leakage (skew=0.02 in Fig 1C top panel) to distributions where most interactions cluster near zero with few positive values when leakage is high (skew=11.2 in Fig 1C bottom panel). What metabolic processes drive this variation?

To answer this, we derive analytical approximations for interaction skewness starting from a simplified consumer–resource model which has no environmental context dependence (i.e., the MacArthur limit with *g*_*i*_(*c*_*α*_) = *R*_*iα*_*c*_*α*_ where interactions do not depend on resource concentrations. See S5) and perturbatively expanding it for small leakage rates. Despite this simplification, the model reproduces the main qualitative trends seen in simulations (Fig 1D). Skewness is lowest when both growth-rate heterogeneity and leakage are low, i.e., among generalists competing for shared resources without cross-feeding. From this baseline, skew increases in two ways. First, without cross-feeding, greater heterogeneity in resource consumption (*R*_*iα*_) increases skew (Fig 1D top). Most species pairs then share few resources, so interactions are usually weak, with only rare strong competitors. Second, increasing leakage (*l*_*i*_) introduces metabolic interdependence (Fig 1D bottom). Leakage both weakens competition (by diverting carbon from growth) and creates positive interactions through cross-feeding. Cross-feeding strength depends on the product of three terms – leakage fraction, stoichiometric matrix, and growth rates. Because these products are mostly small, most positive interactions are weak; only the rare well-matched pairs produce strong ones, shifting the distribution towards zero and increasing skew. These effects interact: leakage amplifies existing heterogeneity in resource consumption. With uniform growth rates, crossfeeding benefits are evenly spread and add little skew. With heterogeneous growth, benefits concentrate in a few well-matched pairs, producing stronger skew (Fig 1C).

In sum, our model and the analytical calculations indicate that “many weak, few strong” interactions emerge from competition and cross-feeding processes, with the degree of skew controlled by both the individual processes and their combined effects.

### Consistent many weak, few strong interactions across empirical datasets

To test whether the patterns observed in our model hold in empirical data, we compared interaction statistics across nine microbial datasets that span different community origins, in which pairwise interaction strengths have been measured or inferred through different approaches (see Methods and Supplementary Table I). The interaction distributions from all these datasets deviate significantly from Gaussian with values clustered near zero, heavy tails containing a few strong interactions, and a consistent negative bias (Figure 2A, deviation from Gaussian: D’Agostino-Pearson *p <* 0.001 for all datasets, an average of 78% values fall within one SD of zero, 1.7% of values fall beyond three SDs from the mean, and all means and medians ≤ 0. See Supplementary Table II for more details).

**FIG. 2.**
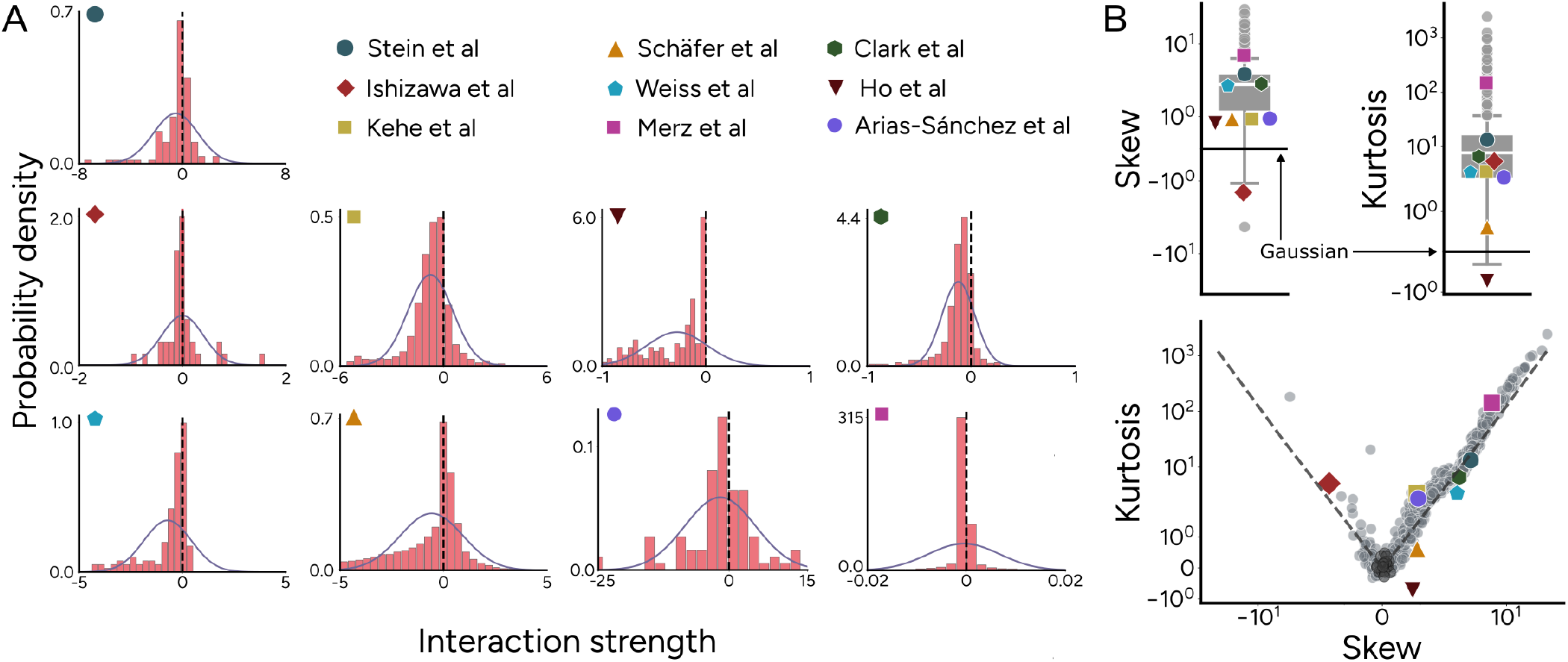
Consistent interaction structure in model and datasets **A)** Interaction distributions from nine microbial datasets from different community origins (Supplementary Table I for more details), showing that all values cluster near zero (dashed line). Solid line indicates a Gaussian with the same mean and variance. **B)** Top: Comparison of statistical measures between model simulations (gray boxes) and empirical datasets (colored points, as in A): Skew (measuring asymmetry and weakness) and excess kurtosis (measuring prevalence of outliers, shortened to kurtosis for brevity). Reference Gaussian values (solid line) are skew = 0 and excess kurtosis = 0. Bottom: Skew plotted against excess kurtosis for both model simulations (gray) and datasets (colored points). Gray dashed line: linear regression fit to model simulations in log-log space. Empirical datasets follow the model scaling with predictive *R*^2^ = 0.78. Black circles indicate communities whose distributions could not be statistically distinguished from a Gaussian (D’Agostino-Pearson *p >* 0.001).

To quantitatively assess whether our model captures this structure, we compared the skewness of the empirical distributions to the distribution of interactions in the 1000 model communities (Fig 1). Despite interactions being measured differently across datasets, skewness, as a dimensionless quantity, captures the overall shape of the distribution, allowing us to compare across different dataset types. Eight of the nine datasets show positive skew, and one shows negative but significant skew, indicating asymmetric distributions consistent with the “many weak, few strong” pattern (Figure 2B, top left). The interactions measured in Ishizawa et al [27] show negative skew, possibly because the dataset contains only seven species (fewer than all other datasets) where a single strong positive interaction shifts the statistic. The model produces a wider spread of skew values than observed in the data (Fig 2B, top left), likely because the randomly sampled metabolic parameters include regimes that generate both near-Gaussian and extremely skewed communities. To quantify the prevalence of outliers beyond skewness, we measured excess kurtosis (see Methods). A Gaussian distribution has zero excess kurtosis and larger values indicate heavier tails. Both model and datasets show substantial excess kurtosis (Fig 2B, top right; median 7.71 vs. 3.36), confirming that the tails contain more extreme values than expected from a Gaussian.

When we plot skew against excess kurtosis (Fig. 2B, bottom panel), the model simulations show a strong scaling relationship in log-log space (linear regression slope = 1.67, *p <* 0.001). This model-derived scaling, with no parameters adjusted to empirical data, gives a predictive *R*^2^ = 0.78 for the skewness–kurtosis values of the nine datasets. The one dataset that deviates the most from the trend is Ho et al [28], possibly because interactions in this study are inferred from a subset of collected metabolomics data (see Methods), which likely ignores many weak interactions, reducing excess kurtosis.

Most model communities were significantly non-Gaussian (98.2 % communities with D’Agostino-Pearson *p <* 0.001). The communities that could not be distinguished from a Gaussian (black circles in Fig 2B bottom panel) cluster near the Gaussian-expected (0, 0) point in the skew-kurtosis space. These communities can occur under specific parameter regimes (e.g., low growth rate heterogeneity), indicating that Gaussian statistics represent a special case within the broader set of distributions our model produces. We tested variations of our model by considering detoxification and pH remediation processes, and still observed skewed, heavy-tailed distributions (S2). The consistency across models, communities, and measurement methods suggests that the non-Gaussian distribution should be the null expectation for metabolite-mediated systems.

### Informed gLV communities better reproduce mechanistic model richness

Having established that metabolite-mediated interactions are non-Gaussian, we next ask whether this matters for predictions of community diversity. If the shape of the interaction distribution carries ecological information, then gLV models parameterized with that shape should outperform those using the conventional Gaussian assumption.

To test this, we constructed what we term “informed” gLV models where interaction statistics were derived from the effective gLV interaction matrices of the mechanistic consumer-resource models (Figure 3A). We generated communities with interactions matrices sampled from three distributions (Figure 3A): First, a Gaussian distribution using the mean and variance. Second, we used a lognormal distribution to capture the skewness of the effective gLV matrices, with many weak and few strong interactions. The lognormal distribution captures the skewness better than alternatives like the gamma distribution, which our analytical calculations suggested (see S3 and S5). And third, we sampled interactions from a correlated lognormal distribution that also incorporates pairwise correlation structure between interactions of the same sign in the effective gLV interaction matrices (see Methods).

**FIG. 3.**
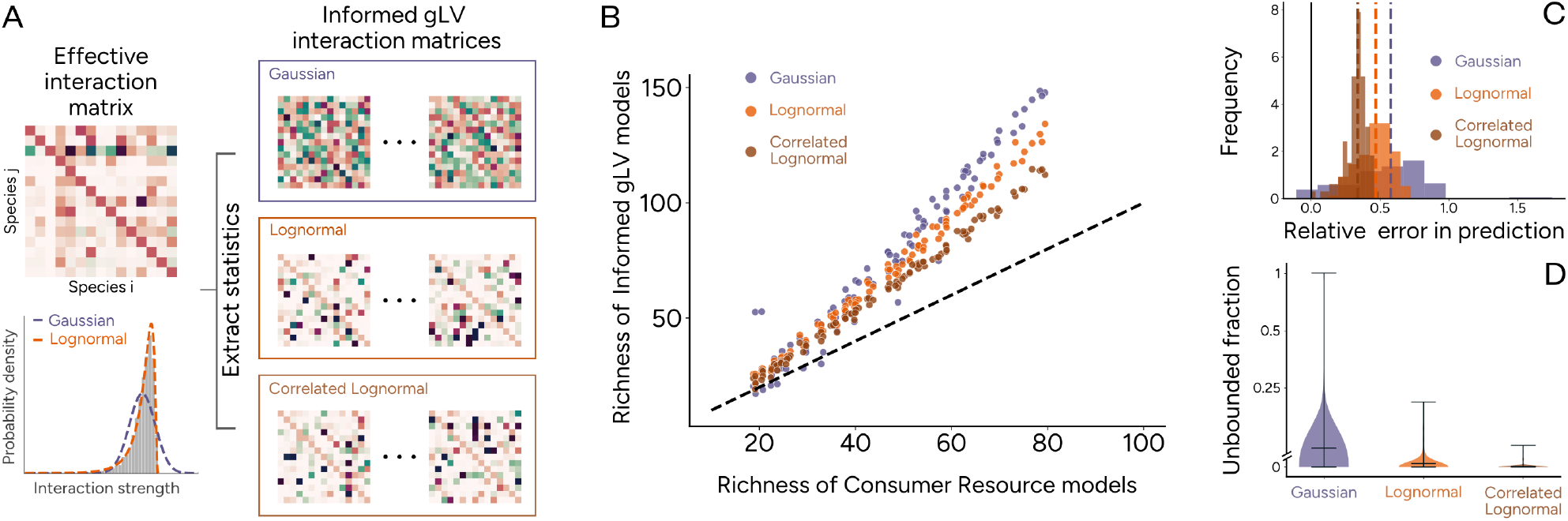
Capturing mechanistic model richness with informed gLV. **A)** Schematic of informed gLV model construction. Statistics are extracted from the effective interaction matrix obtained from consumer-resource simulations, which parameterize a Gaussian, lognormal or correlated lognormal distribution (see Methods), from which multiple interaction matrices are generated. Richness from these informed gLV models is then compared against the original consumer-resource richness. **B)** Community richness as generated by different informed gLV models against mechanistic consumer-resource richness. Dashed line indicates perfect agreement. **C)** Histogram of relative error of log-transformed richness between informed gLV models and consumer-resource model from panel B showing that correlated lognormal has the lowest relative error, while the null Gaussian model has the widest spread. **D)** The fraction of simulations in which abundances grow without bounds in different informed gLV models in B, showing that skewness and correlation aid in model stability.

The error in predicting the final richness of communities simulated using the consumer-resource model grows with increasing richness for all three informed gLV models (linear regression slope between richness and absolute error: Gaussian = 1.08, lognormal = 0.76, correlated-lognormal = 0.48, all *p <* 0.001). The Gaussian model greatly over-predicts richness relative to the mechanistic CR communities it is meant to approximate, while the lognormal and correlated-lognormal models – which incorporate the skewness and the correlations between interactions of the same sign, respectively – track the CR richness more closely (Figure 3C, RMSE normalized by mean ± SD: Gaussian = 0.46 ± 0.21, lognormal = 0.39 ± 0.09, correlated lognormal = 0.29 ± 0.06). Accounting for additional parameters via the Akaike Information Criterion confirms that the correlated lognormal provides the best approximation (AIC: Gaussian = -150.23, lognormal = -179.05, and correlated lognormal = -234.52; lower AIC indicates better fit).

A known limitation of gLV models with Gaussian interactions is that some parameter combinations produce unbounded growth, where abundances increase without limit rather than reaching equilibrium [23]. We find that the fraction of simulations exhibiting unbounded growth decreases as more interaction structure is incorporated (Figure 3D): from 2.6% (Gaussian) to 0.4% (lognormal) to 0.08% (correlated lognormal; Friedman test across parameter regimes: *p* = 0.013). This reduction is consistent with theoretical work showing that correlation structure can suppress unbounded dynamics in gLV systems [33]. Together, these results show that each additional layer of metabolic structure (skewness, then pairwise correlation) improves the ability of phenomenological models to approximate their mechanistic counterparts.

### Experimental diversity is better predicted using lognormally informed interactions

The improvements observed in model-to-model comparisons lead to a stronger test: can lognormally informed gLV models predict diversity in experimental communities better than Gaussian ones? We tested this using the only two datasets out of the nine where (i) time-series experiments involving several species were conducted, from which steady-state diversity could be measured, and (ii) where these experiments were separate from those used to infer interactions between species [27, 28].

The first dataset comes from a synthetic duckweed community of seven species grown in all possible combinations [27]. We inferred interactions from monoculture through four-species communities using the subcommunity-combination method of Maynard et al. [34], reserving larger communities for validation. The second dataset consists of fifteen species from an *in vitro* human gut community [28]. Here, the authors used monoculture spent media metabolomics and pairwise spent media growth experiments to estimate effective resource preferences. The preferences generate a consumer-resource model, which we converted to gLV interactions using the EO framework (see Methods). For each of the two datasets, we calculated the statistics of the interaction distributions (Figure 4A) and used them to generate communities with informed Gaussian and lognormal distributions of gLV interactions. For each model type, we generated 1000 interaction matrices by sampling from the fitted distribution, simulated each to equilibrium, and computed diversity.

**FIG. 4.**
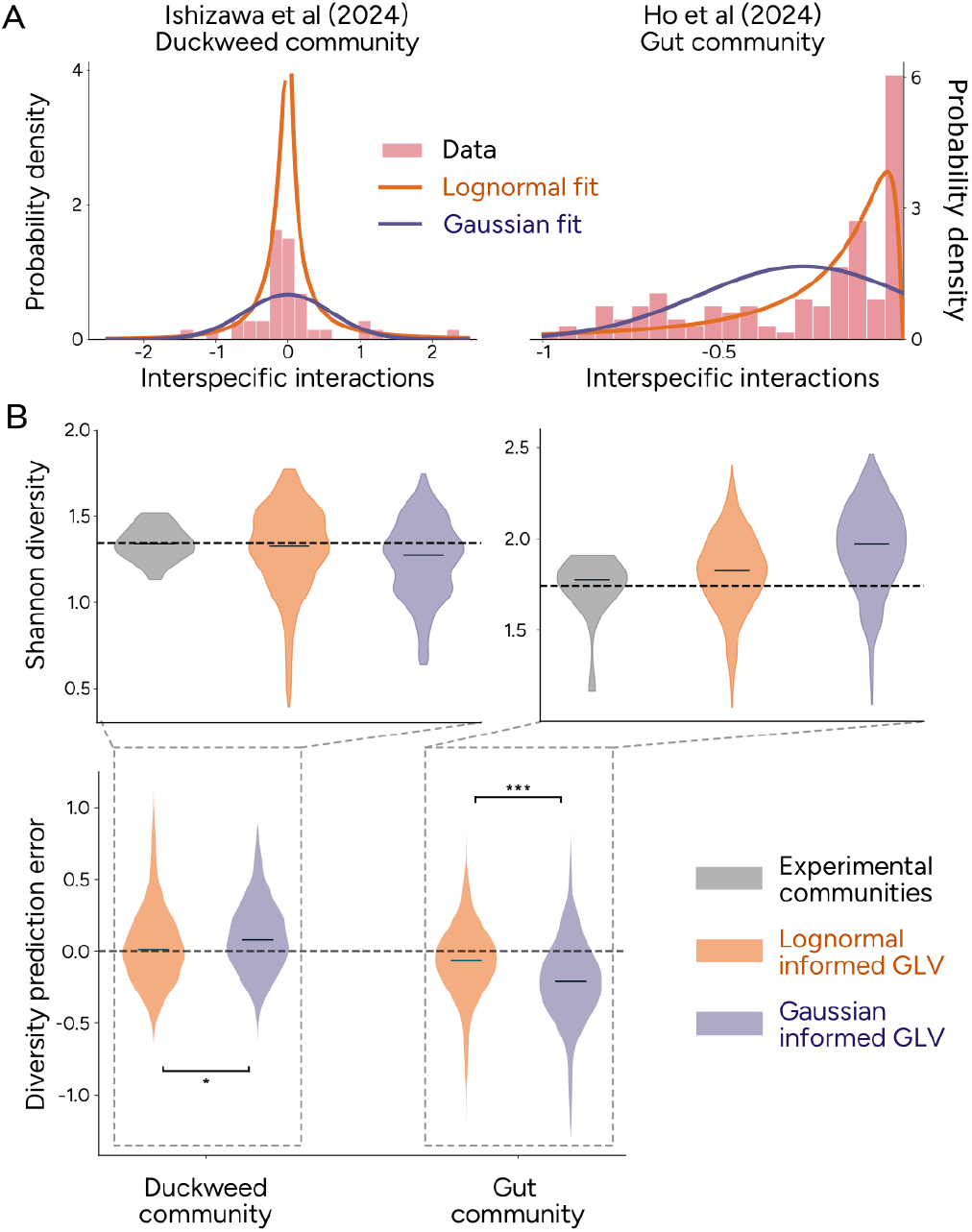
Predictability of experimental communities **A)** Inferred interaction values from Ishizawa et al [27] (duckweed community) and Ho et al [28] (gut community), with lognormal and Gaussian fits. In both datasets, Gaussian approximation fails to capture the empirical distribution **B)** Prediction of Shannon diversity of drop-one-out communities. Upper panels show distribution of predicted diversities from 1000 informed gLV models against a boxplot of experimental values (in gray). Dashed line represents mean diversity of experimental communities. Bottom panel shows the distribution of prediction errors (observed minus predicted diversity) for both datasets and for both models. ^∗^p *<* 0.05, ^∗∗∗^p *<* 0.001, Mann-Whitney U test.

Next, we used the informed Gaussian and lognormal models to predict Shannon diversity of drop-one-out communities (six-species for duckweed, fourteen-species for gut). The lognormally informed gLV model predictions cluster closer to experimental diversity than the Gaussian informed model predictions (Figure 4B, top), showing significantly smaller error (observed versus predicted diversity) for both datasets (Figure 4B, bottom, Mann-Whitney *U* test: *p* = 0.0133 for duckweed, *p <* 0.001 for gut). The improvement is larger in the gut community (mean absolute error: Gaussian = 0.29, lognormal = 0.20), consistent with our simulation results showing that distributional assumptions matter more for larger communities (in Figure 3B). We highlight that in both datasets, the interaction distribution inference was made on smaller communities (up to four species in the case of duckweed and co-cultures in the case of gut), while the prediction was made on a larger community. The consistent improvement of lognormal over Gaussian informed gLV models across the two datasets with different environments and inference methods emphasizes the core finding: metabolic processes generate characteristic non-Gaussian interaction structures, and capturing them is essential to predict diversity.

## III. DISCUSSION

In macroorganisms, patterns of “many weak, few strong” interactions have been repeatedly measured, despite different methods of estimation [12, 35, 36]. These skewed interaction distributions have been attributed to body-size allometry, where interaction strength scales with predator-prey mass ratios [26, 37, 38], or shown to emerge when feasibility and energetic constraints are imposed on trophic food-web models [15], and they are shown to be conserved within taxa [39]. In contrast, these distribution patterns have rarely been explicitly explored in microbes [40], and we would not necessarily expect body-size allometry or trophic food web structure to play a major role. Here we propose that competition for shared resources, together with metabolite exchange through cross-feeding, provides an explanatory mechanism for the same pattern. Strong competitive interactions can arise when resource preferences overlap, with specialization weakening most pairwise effects. For cross-feeding, the mechanism is different. Leakage uniformly weakens competition, while the resulting positive interactions are mostly weak and heterogeneous, depending on the product of multiple independent factors. This effectively shifts the distribution closer to zero and increases skew.

What do these results imply for how theoretical models are parameterized? Since May’s foundational work in the 1970s [8], interactions in gLV models have been drawn from Gaussian distributions, an assumption adopted for mathematical convenience rather than biological realism [23]. Recent theoretical work has shown that the full distribution of interaction strengths, and not just its mean and variance, determines community stability [41, 42], and impacts species abundance distributions [11]. However, without a known target to approximate, these results remain qualitative observations of the behavior of the model rather than quantitative predictions. Consumer-resource models provide one such target. By specifying the underlying metabolic rules, which better capture the underlying processes, we can determine which distributional assumptions best recover community properties like stability and diversity. Our model-to-model comparisons show that each additional layer of distribution structure progressively improves the ability of gLV to approximate consumer-resource richness, and the improvement carries through to experimental communities, where lognormally informed models predict diversity more accurately than Gaussian ones. These improvements arise because in Gaussian models, interactions are symmetrically spread around a mean, implying that moderate-strength interactions are common. In contrast, the “many weak, few strong” structure concentrates probability density near zero, with rare strong outliers. Most species pairs therefore do not affect each other much, and community dynamics may be governed by a smaller effective number of key interactions rather than the full *S*^2^ matrix. This structure not only promotes stability (by reducing unbounded growth in simulations) but is poorly approximated by symmetric distributions. Conceptually, our approach replaces the generic Gaussian prior with one grounded in metabolic constraints, and the progression from Gaussian to lognormal to correlated lognormal mirrors the Bayesian intuition that better-informed priors yield better inference.

This framing has practical implications. Measuring microbial interactions typically involves comparing abundances in coculture versus monoculture, yielding log-ratio metrics [43]. However, these metrics cannot directly parameterize gLV models and hence cannot predict community dynamics. Inferring gLV-compatible interactions requires either time-series data, resource-use information or subcommunity experiments, all more demanding than simple yield comparisons. Even when such inference is possible, the number of pairwise interactions scales as *S*^2^ with community size, making exhaustive measurement infeasible for large communities. Our results suggest an alternative: one can characterize the shape of the interaction distribution from a subset of measurements and use these statistics separately for positive and negative interactions to parameterize a gLV model with skewed interactions. As we show, this approach captures community-level diversity even when individual interactions and exact species abundances are unknown, offering a recipe for making predictions in large community experiments.

The limitations of our study stem from assumptions in the consumer-resource model. Our model considers well-mixed communities with multiple abiotic resources supplied at constant rates, fixed metabolic strategies, and no environmental fluctuations. These assumptions correspond most closely to chemostat experiments with defined multi-resource or rich media. Natural communities, and even many synthetic communities, violate these assumptions in ways that could affect our conclusions about predictability. First, several mechanisms can generate fluctuating interactions even at stationarity: biotic resources [44], diauxic shifts in consumption [45], obligate cross-feeding through auxotrophies [46], or externally driven environmental fluctuations [47]. Our framework assumes that a single, environmentally-dependent interaction distribution characterizes the community. When interactions fluctuate, or the environment changes, whether a previously inferred distribution retains predictive power is an open question. Second, summarizing interactions through distributional statistics assumes that multiple resources are present, as in rich media. When community composition depends strongly on which specific resources each species consumes and leaks, like polymer-breakdown trophic cascades [48], or coexistence on a single supplied resource [18, 20], mean and variance no longer capture the full range of underlying metabolic processes. Lastly, although we have shown that the use of the lognormal distribution can increase predictive power, different distributions might be suitable for different purposes: a Gaussian distribution uses fewer parameters and is easier to infer experimentally in smaller communities, or the gamma distribution may be more analytically tractable for some theoretical contexts.

Despite these constraints, our work connects metabolic mechanisms to community-level outcomes, showing that interaction structure, and not just interaction strength, shapes microbial diversity. That similar distributional patterns arise across ecological domains through distinct mechanisms points toward general constraints on how interactions are organized in diverse communities, and identifying these constraints is a natural next step.

## METHODS

### Consumer-resource models

#### Model assumptions

We consider a model with S number of species and M number of resources, with species i having abundance n_*i*_ and resource α having concentration c_*α*_. Here we list key assumptions of the model, the motivations for choosing them and where they differ from previous studies. We assume resources are carbon sources and hence, need to be externally supplied in a chemostat setup or have to be produced through cross-feeding. Hence, the supply of resources *σ*(c_*α*_) = *δ*(s_*α* −_ c_*α*_) where *δ* is the dilution rate, and s_*α*_ is the supply concentration of the resource. This is different from MacArthur’s original model [17] where resources are biotic in nature and can reproduce by themselves, i.e., resource supply has a logistic form. Next, we describe the per-capita growth rate with a Monod form, motivated by the wide range of observed affinity constants and Monod-like dynamics of batch culture experiments [49, 50]. Hence, the per capita growth rate g_*i*_(c_*α*_) = R_*iα*_c_*α*_/(K + c_*α*_), where R_*iα*_ is the maximal growth rate and K is the resource affinity constant. The consequence of this choice is that it introduces context-dependence, i.e., interactions at steady state are dependent on the resource concentrations. Lastly, we consider a conservation of energy in resource cross-feeding, which is a natural assumption that more carbon atoms cannot be produced than what is taken up. We implement this by drawing the stoichiometric matrix from a uniform distribution *U*(0, 1/(*M* − 1)) and setting the diagonal element D_*αα*_ = 0. This ensures that the column sum is always less than unity, and hence, no more resources can be produced than taken up, similar to previous studies [18].

Analytical solutions of steady state abundances and concentration are not possible with the Monod per capita growth rate and the chemostat-like supply. Hence, in Fig 1D and S5, for our analytical calculations of the interaction skew, we consider a simplified version with g_*i*_(c_*α*_) = R_*iα*_c_*α*_ and *σ*_*α*_ = c_*α*_(s_*α* −_ c_*α*_), which reduces to MacArthur’s original model [17].

### Microbial interaction datasets and comparison

We use a total of nine datasets where interactions have been inferred through different methods. See Supplementary Table I for a description of all nine datasets. Seven datasets had reported interaction values: Stein et al [51], Kehe et al [4], Schafer et al [40], Weiss et al [52], Clark et al [53], Merz et al [54], and Arias-Sánchez et al [55]. The other two datasets included abundance or metabolomics data, which we used to infer effective gLV interactions as outlined below.

#### Synthetic duckweed community [27]

The community consists of seven species grown in duckweed fronds for 10 days. The experiment included all combinations of the species, ranging from monocultures to all seven grown together. We applied the data to the method described by Maynard et al [34] where they minimize the difference between predicted abundance (from inverting a gLV interaction matrix) and observed abundance, across different species combinations. We obtained the interaction matrix from combinations of 1-, 2-, 3-, and 4-species. For predicting diversity using informed interaction matrices later, we use the six-species (drop-one-out) communities, which were not used to quantify the interactions.

#### In vitro human gut community

This community consists of 15 species, where authors analyze the spent media from monocultures and pairwise combinations. Looking at metabolomic profiles before and after culturing, they infer effective resource groups in rich media and conclude that most interactions are competitive. They parameterized a resource preference matrix by grouping the used metabolomic profiles into effective resource groups, finding that 25 different effective resource groups best describe observed abundances. We take this matrix as the basis for the metabolic processes that occur in the community. Three species (*Bt, Pd, Csc*) were inferred to be pure specialists, that is, they consumed only their unique resource group. Obtaining the interaction matrix with these species will result in them not interacting with any other species, effectively being independent. Hence, we discard them for the informed gLV models. But we note that their non-interaction with the community is a result of the inference procedure where weak overlap in resource groups might not be captured. Hence, the excluded species could still have an effect on community diversity. Therefore, we include them in the diversity predictions, even though, by excluding them, our informed gLV models are likely underestimating the weak interactions. Even after excluding them, the interaction matrix is quite sparse (approx. 28%) because of some non-overlapping resource groups (see S6). Hence, we randomly add the sparsity on top of our generated gLV matrix (for both Gaussian and lognormal interactions) by multiplying a random matrix filled with zeros and ones corresponding to the observed sparsity. These simulations are then compared to 14-species (drop-one-out) communities.

#### Measured distribution statistics

Previous literature on gLV models considers interaction matrices such that the diagonal elements representing the strength of intraspecific limitation have a positive value. We flip the sign of the interaction strengths from datasets to align with this convention, which also helps in comparing to the interaction values from our model. We measure two dimensionless quantities from both the model and all nine datasets. First, we compute skewness, defined as the ratio of the third cumulant (see below) to the cube of standard deviation [56]. This quantity captures whether the mean of the distribution is left (negative skew) or right (positive skew) of the median, thereby indicating how symmetric the distribution is. Then, we compute the excess kurtosis, which is similar to skew, except, with the fourth cumulant and the fourth power of standard deviation subtracted by three [13, 56]. This measure is used to examine whether there are outliers in the distribution, with larger values indicating stronger outliers. The definition for the skew and kurtosis of a set of values {x_1_, ·I ·I ·I, x_*N*_ } with mean 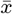 is given by

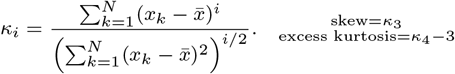

The excess kurtosis is subtracted by three to account for a Gaussian distribution which would have kurtosis of 3 otherwise. With this, a Gaussian has zero skew and zero excess kurtosis.

### Effective gLV interactions

#### Converting consumer-resource to Glv

We apply the Environment-Organism (EO) framework to convert metabolic processes into interactions [25]. As a reminder, given a general consumer-resource model with abundance 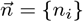 and resource concentrations 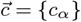

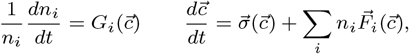

effective instantaneous interactions are given by 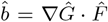 and effective growth rate 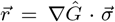, where *Ĝ* and 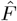 are the matrices corresponding to the respective collection of sensitivity and impact vectors evaluated at the steady state resource concentrations, and the gradient ∇ is with respect to the resources. These matrices correspond to the stationary equilibrium solution given by 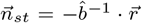. This means that a gLV model initialized with the effective interactions and growth rates will result in the same equilibrium abundance as observed in the consumer-resource model. However, since these are instantaneous interactions, they do not strictly correspond to gLV interactions. To circumvent this issue, we normalize the effective interactions by the growth rates and the self-interaction. In doing so, all intraspecific interactions become unity, i.e., *a*_*ii*_ = 1 and this now corresponds to a gLV model whose dynamics are given by

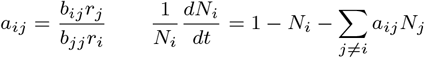

with *N*_*j*_ = *n*_*j*_ *r*_*j*_/*b*_*jj*_ . This allows us to make comparisons to the standard gLV and use statistics of *â* (the matrix of interspecies interactions) to generate informed models.

#### Informed gLV models

Given an effective interspecies interaction matrix *â*, we construct informed gLV models at three levels of detail, each adding one layer of structure from the effective interactions.

##### Gaussian informed model

The simplest model draws all off-diagonal interactions from a single normal distribution *N* (⟨*a*⟩, *σ*^2^(*a*)), where ⟨*a*⟩ and *σ*^2^(*a*) are the mean and variance of the off-diagonal entries of *â*.

##### Lognormal informed model

Because the effective interactions are skewed and contain both positive and negative values, we fit separate lognormal distributions to each sign class. We partition the offdiagonal entries of *â* into positive entries *a*^+^ and negative entries *a*^−^. For the negative values, we work with their magnitudes | *a*^−^ | so that the log transform is well-defined. We then fit normal distributions to the log-transformed values of each sign class:

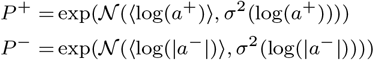

where the negative sign in *P*^−^ restores the original sign after exponentiation. To generate an informed interaction matrix, each off-diagonal entry *a*_*ij*_ is drawn from *P*^+^ with probability m or from *P*^−^ with probability 1 − m, where m is the fraction of positive values in *â*.

##### Correlated lognormal informed model

The lognormal model treats each entry *a*_*ij*_ independently. However, in the effective interaction matrices, pairs of interactions of the same sign – where both *a*_*ij*_ and *a*_*ji*_ are positive, or both are negative – are correlated. To capture this, we replace the independent normal distributions used above with bivariate normals. For positive-positive pairs, the log-transformed values 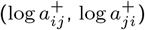 are drawn from a bivariate normal with the same marginal means and variances as in the lognormal model, but with covariance 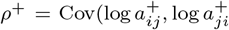 Cov(log The analogous covariance ρ^−^ is computed for negative-negative pairs. Entries are then exponentiated and assigned signs as before. Pairs where *a*_*ij*_ and *a*_*ji*_ have opposite signs are drawn independently from the appropriate marginal distributions.

#### Simulation details

##### General settings

All simulations were performed in Python using the LSODA algorithm [57] from the package numbalsoda v0.3.5. Consumer-resource models were initialized with all species abundances at 0.1 a.u. and resource concentrations at their supply values. Generalized Lotka-Volterra models were initialized with all species abundances at 0.1 a.u. Both model types were integrated until T = 10000, which was sufficient for all simulations to reach stationarity. A species was considered extinct if its final abundance fell below 10^−4^. For gLV models, a simulation was classified as exhibiting unbounded growth if any species abundance exceeded 10^8^.

##### Consumer-resource model parameters

The Monod affinity constant was set to K = 1 and the dilution rate to *δ* = 0.2 for all simulations. Growth rate matrices R_*iα*_ were drawn from a lognormal distribution (alternative distributions also produce skewed interactions; see S2). Leakage fractions l_*i*_ were drawn from a uniform distribution U(l_*min*_, l_*max*_), allowing control over the mean and variance of leakage and thereby the extent of cross-feeding. The stoichiometric matrix *D*_*αβ*_ was drawn from *U*(0, 1/(*M* − 1)) with *D*_*αα*_ = 0, ensuring that no more resources are produced than consumed.

##### Computing effective interactions

After each consumer-resource simulation reached stationarity, we applied the EO framework to obtain effective gLV interactions. The gradient ∇ Ĝ and 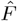 were computed analytically from the Monod functional form and evaluated at the stationary resource concentrations. This procedure requires that the number of surviving species does not exceed the number of resources (see S1); this condition was satisfied in all reported simulations because competitive exclusion ensures that at most M species coexist.

##### Informed gLV model parameters

For all informed gLV models (Gaussian, lognormal, and correlated lognormal), diagonal elements were set to unity (a_*ii*_ = 1), matching the normalization of the effective interaction matrices. Off-diagonal elements were drawn as described above (see Informed gLV models). The fraction of simulations exhibiting unbounded growth under each distributional assumption is reported in Fig 3D.

##### Richness prediction and model comparison (Fig 3)

For each of 100 parameter combinations (varying species number S ∈ [60, 220], resource ratio M/S ∈ [1, 2], and mean leakage ∈ [0.3, 0.6]), we simulated 50 consumer-resource communities sharing the same growth rate statistics but with independently drawn consumption and stoichiometric matrices. The interaction statistics from these 50 communities were used to parameterize informed gLV models, from which 100 interaction matrices were generated for each of the three distribution types (Gaussian, lognormal, correlated lognormal). This yielded 5000 consumer-resource and 30000 gLV simulations in total. Growth rate matrices were drawn from a lognormal distribution with mean 1/M and standard deviation 10/M. Of the M total resources, half were randomly chosen to be externally supplied. The remaining resources could only be produced through cross-feeding.

## Acknowledgments

We thank the members of the Mitri lab, Oliver Meacock, Vit Piskovsky, Ming Liu, Sandro Azaele, and Kevin Foster for helpful feedback. We also thank Po-Yi Ho for pointers on their dataset and Daniel Maynard et al for wonderfully reproducible and documented code from their paper. This work was supported by the Swiss National Science Foundation (SNSF) through Project grant 320030-236360, the National Center of Competence in Research (NCCR) Microbiomes (grant number SNF 51NF40 180575), and the Faculty of Biology and Medicine at the University of Lausanne.

## Code and Data Availability

Datasets used from previously published papers can be accessed at their original published location. The post processed interaction distributions, and all other codes can be found at https://github.com/Mitri-lab/interaction_distributions.

