## Supplementary Information for "Metabolic processes shape microbial interaction distributions"

### Supplementary Information
#### Metabolic processes shape microbial interaction distributions

Prajwal Padmanabha and Sara Mitri
*Department of Fundamental Microbiology, University of Lausanne*

| Paper | Species richness | Community Origin | Description |
|---|---|---|---|
| Stein et al [1] | 11 | Mouse gut (*in vivo*) | Interactions inferred by fitting a gLV model to time-series grouped relative abundance data at genus or family level. |
| Weiss et al [2] | 12 | Mouse gut | Dataset reported ratio of coculture growth to monoculture growth of a synthetic community that is representative of mouse gut microbiome. Interactions were computed by log-transforming the reported values. Despite common origin to Stein et al [1], the interaction inference method is different and this dataset contains specific strains. |
| Kehe et al [3] | 20 | Soil | Interactions measured using log ratio of coculture abundance to monoculture abundance across 40 different carbon source environments in kChip |
| Schäfer et al [4] | 224 | Plant leaf | Interactions computed using biomass production rate alone or in the presence of a partner from genome-scale metabolic models of bacterial strains isolated from *Arabidopsis thaliana* leaves. |
| Ishizawa et al. [5] | 7 | Duckweed | Data contains abundances across all possible species combinations grown on duckweed fronds (monoculture through seven-species). Interactions were inferred by us using the subcommunity method of Maynard et al. [6] by using monoculture through four-species communities. |
| Ho et al [7] | 15 | Human gut | Data contains fresh media and spent media metabolomics of monocultures grown on BHI media which is used to infer effective resource groups and growth rates. Interactions were inferred by us using the growth rates on effective resource groups and the EO framework. The effective resource group computation results in discarding nearly 90% of metabolomic features that contribute weak effect. This results in interactions being computed as being zero while in reality, they can be weakly negative or positive. This spike at zero impacts the kurtosis value, decreasing it from the model trend. |
| Clark et al [8] | 25 | Human gut | Dataset already contains interactions obtained using Bayesian inference from time-series abundance data. Despite common origin to [7] and both being synthetic communities, the inference methods of interaction is different. Additionally, Clark et al [8] choose strains to optimize for butyrate production while Ho et al use a community that is represenative of the human gut microbiome. |
| Arias-Sánchez et al [9] | 11 | Metal Working Fluid | Dataset contained interactions measured from a metal-working fluid community that was subjected to artificial selection experiment. Interactions were reported as $\log_2$ of ratio of yield in coculture vs monoculture. We took reported interactions only from the ancestral strains to avoid influence of selection experiment. |
| Merz et al [10] | 162 | Marine | Dataset reported interactions (measured as Jacobian elements fitted using empirical dynamic modeling) in a six-year time series data of coastal marine community. |

TABLE I. Description of microbial datasets used in the paper

| Paper | Mean strength | Median strength | % -ve values | values $<$ 1 SD from 0 | values $>$ 3 SD from mean | Skewness | Excess Kurtosis |
|---|---|---|---|---|---|---|---|
| Stein et al | -0.518 | -0.084 | 59% | 85.45% | 1.82% | 3 | 13.33 |
| Weiss et al | -0.722 | -0.259 | 80.91% | 79.39% | 2.29% | 1.96 | 3.36 |
| Kehe et al | -0.765 | -0.615 | 80.08% | 75.02% | 3.08% | 0.91 | 3.36 |
| Schäfer et al | -0.576 | -0.005 | 55.2% | 73.88% | 0.02% | 0.92 | 0.61 |
| Ishizawa et al | -0.03 | -0.07 | 59.5% | 80.95% | 2.38% | -1.39 | 5.09 |
| Ho et al | -0.278 | -0.152 | 100% | 65.15% | 0.0% | 0.8 | -0.72 |
| Clark et al | -0.122 | -0.093 | 89% | 73.33% | 2.17% | 2.05 | 6.57 |
| Arias-Sánchez et al | -1.68 | -0.98 | 61.29% | 77.42% | 2.15% | 0.94 | 2.73 |
| Merz et al | -0.004 | 0 | 69.07% | 93.15% | 2.02% | 6.10 | 142.76 |
| Gaussian | NA arbitrary | Same as mean | NA depends on mean | NA depends on mean | 0.14% | 0 | 0 |

TABLE II. Interaction distribution statistics from datasets used in the paper and comparison to an arbitrary Gaussian. Ho et al [7] consider only the most relevant metabolomic features leading to many zero values, while Scäffer et al [4] ignore interactions whose strengths are $> 5$, both leading to fewer outliers.

#### 1. ENVIRONMENT-ORGANISM INTERACTIONS VS TIMESCALE SEPARATED INTERACTIONS

We show in this section that different methods of obtaining effective interactions from a consumer resource model have different consequences, statistically speaking. We begin by considering a generic consumer-resource model where the abundance of species $i$ is given by $n_i$ and the concentration of resource $\alpha$ by $c_\alpha$

$$\frac{1}{n_i}\frac{dn_i}{dt} = G_i(\vec{c}) \qquad \frac{d\vec{c}}{dt} = \vec{\sigma}(\vec{c}) + \sum_i n_i \vec{F}_i(\vec{c}), \tag{1}$$

where $F_i$ and $G_i$ are the sensitivity and impact. Example dynamics for a set of species and resources are shown in Fig 1A. The EO interactions and growth rates are given by

$$a_{ij}^{EO} = (\nabla G \cdot F)_{ij} \qquad r_i^{EO} = (\nabla G \cdot \sigma)_i \tag{2}$$

This comes from the stationary state of an exact integro-differential equation that takes into account the the feedback between organisms and the environment when integrating out the resources [11]. An alternate way to obtain interactions is to consider that the dynamics of resources is much quicker than that of species – an assumption that does not hold true in the chemostat [12] - and then to calculate the equivalent quasi-stationary resource concentration to be input in the species dynamics. This timescale separation trick is quite common and was originally originally used by MacArthur to derive interactions in a biotic supply model (which we shall return to later) [13]. In fact, many studies have independently converged on the same interaction matrix calculation following this technique [14–17]. For the generic CR model defined above, the derivation is more involved, as presented initially in the supplementary information of Koffel et al [14]. The trick involves defining a new matrix that effectively captures resource-resource interactions. This matrix $H$ is given by

$$H_{\alpha\beta} = \sum_{k,\gamma} \frac{\partial F_{k\alpha}}{\partial c_\beta} F_{k\gamma}^{-1} \sigma_\gamma - \frac{\partial \sigma_\alpha}{\partial c_\beta} \tag{3}$$

where the matrices and vectors are evaluated at resource concentrations such that $\dot{\vec{c}} = 0$, and $F^{-1}$ represents the pseudo-inverse. Given this resource-resource interaction matrix, the timescale separated growth rates and interactions are given by

$$a_{ij}^{TS} = (\nabla G \cdot H^{-1} \cdot F)_{ij} \qquad r_i^{TS} = (\nabla G \cdot H^{-1} \cdot \sigma)_i. \tag{4}$$

Note that this is very similar to the EO framework (Eq. 4), albeit not coming from a strictly valid approximation in the case of chemostat dynamics. Despite this, due to the commonality of the structure and the evaluation of both frameworks at stationarity, they produce the same estimated populations if $-a^{-1} \cdot r$ is considered. gLV simulations using either interaction matrix results in equilibrium abundances that correspond to the consumer-resource simulations, as long as the number of resources in the system $M$ is greater than the number of species $S$. The moment this condition fails, these interaction matrices do not provide a good estimate of the stationary abundance, a fact that possibly underlies the results in [17] where

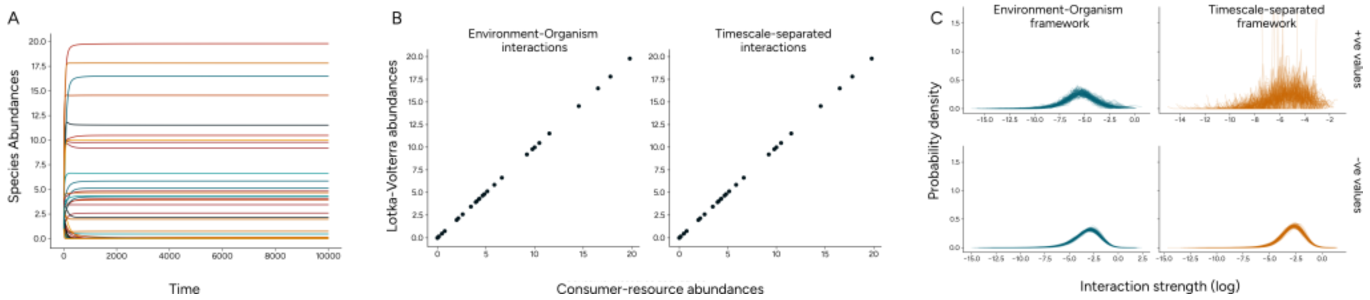


FIG. 1. Applicability of Environment-Organism framework as opposed to timescale separation approximation **A)** Dynamics of population abundances from consumer-resource model with competition and cross-feeding, starting from S = 100 species and M = 100 resources **B)** Comparison between observed abundance (from CR models) and abundances from effective gLV models. The gLV interactions are directly taken (not converted into statistics like with informed models) **C)** Distribution of interactions from the two frameworks. Top panels show positive interactions on a log scale and bottom panels show negative interactions on a log scale

their estimation from the interaction matrix almost never coincides with the consumer-resource abundances. We show the exact prediction in Fig. 1B where the $x$-axis represents the abundance from the consumer-resource simulation and the $y$-axis is the abundance given by $-a^{-1} \cdot r$ of the appropriate model.

However, we use the EO framework in our work to avoid the underlying assumption of timescale-separation. Considering the interaction matrix through timescale-separation leads to the inverting of $H$ matrix, for which we need to consider a pseudo-inverse of $F$. These two inversions lead to unreliable interaction distributions – despite starting with the same growth rate and cross-feeding statistics, different communities give rise to different interaction distributions as shown in Fig. 1C. The EO framework yields more consistent distributions, plotted on a log-scale for both positive and negative interactions. However, the timescale-separation framework produces a different distribution in each simulation, especially visible in the positive interactions subplot (CV of mean interaction strength: EO framework = 0.17, TS framework = 0.30). This means that making the timescale separation assumption would generate less consistent informed interaction statistics. Hence, parameterizing an informed gLV model might lead to unreliable estimates of community richness or diversity.

This result might arise from the process of taking statistics *after* performing a mathematical operation on a set of random variables. In particular, random matrices that have consistent statistics before inversion can result in very different statistics post inversion, due to eigenvalues close to zero blowing up after inverting [18]. The presence of such non-linearities through the pseudo-inverse in timescale separated interactions implies that the obtained statistics might not consistently follow the same distribution. To avoid these issues, we utilize the exact EO framework in our paper.

#### 2. SKEWED INTERACTIONS IN MODEL VARIATIONS

A critical question is whether the skewed interaction is based on model assumptions (of drawing underlying distributions) or the nature of cross-feeding and positive interactions. Here, we present three variations all of which show skewed interaction distributions.

**Varying growth rates** We first look at whether changing growth rate distributions can impact interaction skew. We simulate communities with growth rates having the same mean and variance, but from either a lognormal or gamma distribution. We consider 200 species and 200 resource for this test with leakage rates between 0 and 0.4. Figure 2A shows that regardless of the growth rate distribution, we observe the skew (Gamma growth rates = 8.01, Lognormal growth rates = 3.13 compared to Gaussian expectation of zero). A note is that for the chosen values of mean and variance that we use in the main text, we cannot obtain a uniform distribution (since the minimum value necessary would be negative). However, for feasible values (up to growth rate CV around 5), we still see the skew (around 5.76), indicating that interactions arise from niche overlap and not the particular distribution of growth rates.

For extending to other models, we describe two changes here: one in which we relax and modify the assumptions used in our model while adding detoxification, and the second in which we describe a new mechanism of positive interaction through pH mediation.

**Crossfeeding-detoxification model** In this extension of the model, we relax certain assumptions made in the original model. Previously, we considered the Monod affinities for growth to be constant. Here, we sample the affinities ($K_{i\alpha}$) from a lognormal distribution of positive values (with mean $= 1$ as before and sd $= 2$). While previously the stoichiometric matrix was drawn from a uniform distribution, here we draw it from a lognormal distribution (with mean = 0.1, sd = 1) and normalize the row sum to one. This allows for non-uniformity in the cross-feeding chains but conserves total carbon content and reflects the possibility of specific conversion pathways. Finally, we were modeling co-utilization of resources by dividing the total growth rate by number of resources consumed. Here, we modify that to consider a proteome-allocation tradeoff such that the total growth rate is normalized to unity. We model detoxification as an investment of resources taken up, with a fraction $f_i$ by species $i$ as shown in Fig 2B left panel. The presence of toxins adds an additional death rate to species, and all species degrade toxins at the same rate through dynamics similar to that of resources, i.e., toxins are degraded whenever a resource is used for growth.

With these modifications to the model in the main text, the dynamics for abundance $n_i$ and resource $c_\alpha$ is given by

$$\frac{1}{n_i}\frac{dn_i}{dt} = \sum_\alpha (1-l_i)(1-f_i)\frac{R_{i\alpha}c_\alpha}{K_{i\alpha}+c_\alpha} - \sum_x \frac{\eta_{ix}t_x}{Q_{ix}+t_x} - \delta \quad (5)$$

$$\frac{dc_\alpha}{dt} = \delta\left(s_\alpha - c_\alpha\right) - \sum_i n_i \frac{R_{i\alpha}c_\alpha}{K_{i\alpha}+c_\alpha} + \sum_i l_i n_i \sum_\beta D_{\alpha\beta}\frac{R_{i\beta}c_\beta}{K_{i\beta}+c_\beta} \quad (6)$$

$$\frac{dt_x}{dt} = \delta(s_x - t_x) - \gamma t_x \sum_i (1-l_i) f_i n_i \sum_\alpha \frac{R_{i\alpha}c_\alpha}{K_{i\alpha}+c_\alpha} \quad (7)$$

where $t_x$ now represents the concentration of a toxin $x$, $\eta_{ix}$ the death rate due to the toxin, $Q_{ix}$ is the half saturation constant of the effect of toxin, $\gamma$ is the degradation rate of the toxin with supply concentration $s_x$, and all other parameters are the same as the model in main text. Figure 2B middle and right panels show that these interactions observed at stationarity also display large skew values (skewness = 17.76), here plotted as log transformed positive and negative interactions.

**pH mediation model** We describe a different source of positive interactions, now driven only by competition and pH mediation. This is motivated by the impact of resource concentrations on the pH of the medium and, subsequently, on the growth rates. We modify the per capita growth rate in the competitive consumer-resource model compared to the Monod functional form, as shown in Figure 2C left panel. Here, increased resource concentrations lead to lower growth rates, leading to an optimal concentration for maximal growth. However, if resource concentration is suboptimal for growth, reducing it increases the per capita growth rate, leading to positive interactions. To observe positive interactions in this model, our steady state resource concentration must be in the suboptimal regime. Hence, we consider a high dilution rate, close to the community washout ($\delta = 2$ as opposed to 0.2 from before). We observe a skewed interaction distributions as shown in Fig 2C middle and right panels (with skewness = 2.40). Figure 2C middle panel shows fewer positive interactions due to the nature of the model - the dilution rate must be at the precise point to avoid complete washout while also allowing positive interactions to emerge, leading to fewer values.

#### 3. INTERACTION DISTRIBUTION FITTING

Given that we observe skewed distributions, the next question is how to parametrize it. Two standard models that are known to show skewed structures are lognormal and gamma. Typically, it is difficult to tell one distribution from the other. We fit a lognormal distribution as outlined in the Methods. For the gamma approximation, we compute the best fit of the cumulative distribution function (CDF). Given a shape $(\alpha)$ and scale $(\theta)$ parameter, the CDF of a gamma distribution is given by $F(x) = \gamma(\alpha, x/\theta)/\Gamma(\alpha)$, where $\gamma(\alpha, x/\theta)$ is the lower incomplete gamma function and $\Gamma(\alpha)$ is the ordinary gamma function. We fit $F(x)$ to the cumulative distribution of effective interaction values to obtain the gamma distribution parameters.

Figure 3A shows that both distributions fit well the interactions inferred from the CR model (Goodness of fit $R^2$: gamma = 0.95, lognormal = 0.94). However, when we look only at strong outliers (interactions stronger than one standard deviation from the mean), lognormal distribution outperforms the gamma ($R^2$ with outliers: gamma = 0.69, lognormal = 0.90). We proceed to test how well do informed gLV models from these distributions compare against consumer-resource communities. The gamma distributed interactions overpredict the richness more than the lognormally distributed interactions (from 100 simulated communities in Fig 3B: Richness in communities - CR = 32.1 ± 2.98, lognormal = 45.9 ± 4.17, gamma = 79.11 ± 3.91). Even though the predictions of the lognormally distributed interaction model are not particularly accurate, predicted richness is still closer to that generated by the CR model than the gamma distribution. Hence, in the main text, we use the lognormal distribution as an approximation for the effective interactions. This also comes with the advantage that measuring and implementing correlations between log-transformed interactions are easier in the lognormal distribution than the gamma distribution.

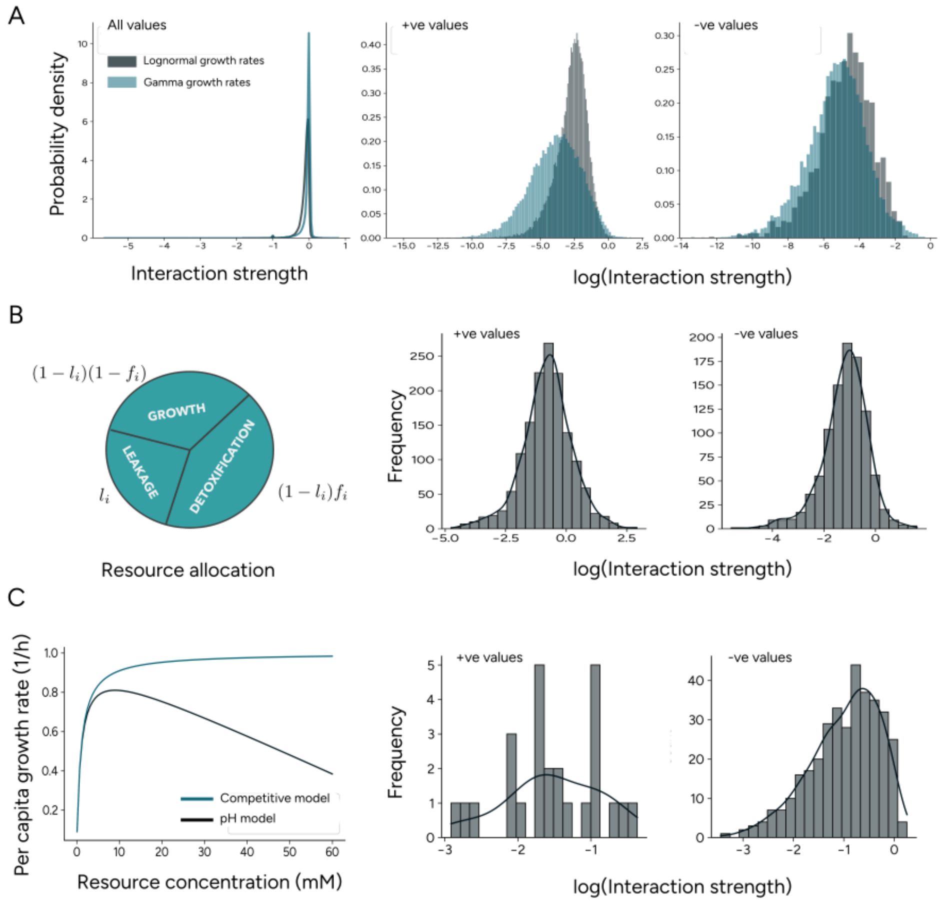


FIG. 2. Different assumptions and models demonstrating consistent interaction patterns **A)** Variation in distribution from which species growth rates are drawn from. Gray bars represent underlying lognormal distribution while blue bars represent underlying gamma distribution. Interactions are plotted separately in the second and third panel on a log scale **B)** Crossfeeding-detoxification model with variable resource affinities. Total resource taken up is split into growth, leakage, and detoxification and shown in the left panel. Log of interaction strengths are shown in the other two panels, with the black line plotted from automatic kernel density estimation **C)** pH remediation model where excessive resource concentration decreases per capita growth rate as shown in left panel. Positive interactions are fewer in this case but still present. Negative interactions still show significant skew compared to a Gaussian

#### 4. SIMULATION PARAMETER RANGES

**Predictability and model stability data** For data in Fig 3B of the main text, we systematically vary number of species, resources and leakage fractions rather than sampling them randomly so that our communities are always within the range in which EO framework is applicable (see S1). We draw growth rate matrices from a lognormal distribution with mean $1/M$ and standard deviation $10/M$. We vary the number of species between $S \in [60, 220]$ in jumps of $40$. For each number of species, the number of resources is chosen as $M = \gamma S$, where $\gamma \in [1, 2]$ in steps of $0.25$. Of these total $M$ resources, we externally supply half randomly chosen ones. For each of these 25 conditions, we consider four sets of leakage fractions. The mean leakage varies between $[0.3, 0.6]$ in steps of $0.1$ - giving a total of 100 conditions. For each condition, we consider 50 consumer-resource communities such that they all have the same mean and standard deviation of growth rates but different consumption matrices. These 50 communities are used to calculate the statistics of interaction matrices used to inform gLV models. Then, we simulate 100 informed gLV models with three possible distribution choices as listed in the Methods section. This results in a total of 5000 CR communities and 30000 gLV communities being simulated to compute the error between consumer resource richness and gLV richness.

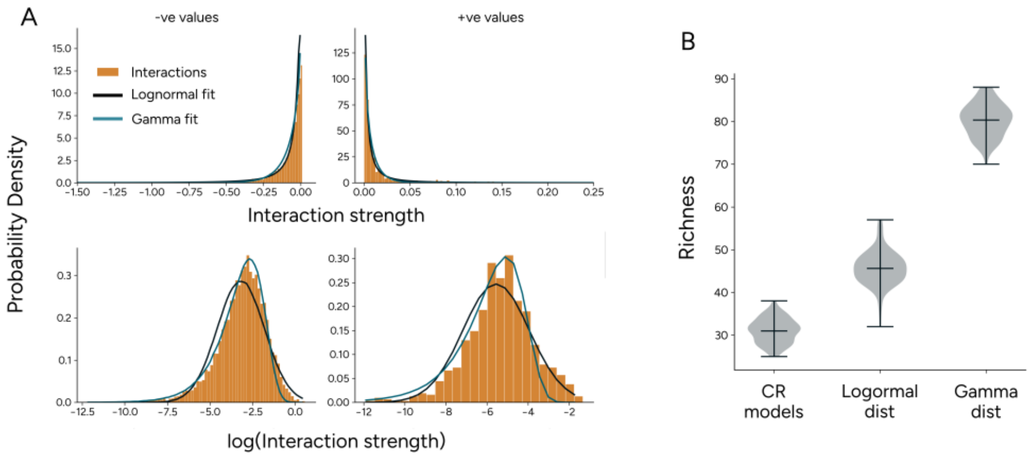


FIG. 3. Fitting interaction distributions to different functions **A)** Approximating effective interactions (orange) from a single consumer-resource community by lognormal (black) and gamma (blue) distributions. Top row shows the direct distribution while the bottom row shows the same plot in log transformed values. Accurate representation of median values by gamma distribution and extrema values by lognormal distribution is evident in the bottom row **B)** Using the statistics from each approximation into an informed gLV model and comparing the predicted richness to the observed consumer resource richness. The lognormal distribution's prediction is closer to the CR models.

#### 5. ANALYTICAL SKEW CALCULATIONS IN MACARTHUR MODEL

In this section, we compute whether a simplified consumer-resource model can analytically capture the skew we observe in a more complex model. The competitive MacArthur model for $S$ species and $M$ resources is defined by the following equations:

$$\dot{n}_i = n_i \left( \sum_\alpha R_{i\alpha} c_\alpha - \delta_i \right) \tag{8}$$

$$\dot{c}_\alpha = c_\alpha \left( s_\alpha - c_\alpha \right) - \sum_i R_{i\alpha} c_\alpha n_i \tag{9}$$

where we assume that the yield of each resource is equal. In quasi-stationarity, i.e., when resource dynamics are much faster than species dynamics, the quasi-equilibrium resource concentrations are nonzero, given by $c^*_\alpha = s_\alpha - \sum_i n_i R_{i\alpha}$. When this is substituted into the equation for the species, we obtain

$$\dot{n}_i = n_i \left( \sum_\alpha s_\alpha R_{i\alpha} - \delta_i - \sum_j \left( \sum_\alpha R_{i\alpha} R_{j\alpha} \right) n_j \right) \tag{10}$$

allowing us to identify the equivalent gLV growth rate and interaction as $r_i = \sum_\alpha s_\alpha R_{i\alpha} - \delta_i$ and $a_{ij} = \sum_\alpha R_{i\alpha} R_{j\alpha}$. Note that we adopt timescale separation here purely to obtain a tractable analytical scaling; unlike the distribution-level estimates of the main text (S1), this qualitative derivation does not require consistency of the interaction statistics across communities, so the inversion pathologies discussed in S1 do not apply. Assuming $s_\alpha = s$ and $\delta_i = \delta$, and noticing that $a_{jj} = \sum_\alpha R^2_{j\alpha}$, we can investigate the statistics of a rescaled interaction matrix as that of $a_{ij}$, up to the scaling imposed by the second moment of distribution of the growth rates. Let us assume that the growth rates of the species on different resources are independently drawn from a single distribution given by $P_R(R)$, where we denote the $m-th$ moment of this distribution by $\mu_m(R)$ and $m$-th cumulant by $\kappa_m(R)$.

Then, the mean interspecies interaction is given by

$$\langle a_{ij} \rangle = \frac{1}{S(S-1)} \sum_{i \neq j} \sum_\alpha R_{i\alpha} R_{j\alpha} = \frac{1}{S(S-1)} \sum_\alpha \sum_{i \neq j} R_{i\alpha} R_{j\alpha} = M \mu_1(R)^2 \tag{11}$$

where we switch summations due to the independence of random variables and use the fact that $i \neq j$ leads to uncorrelated variables. For the variance, we first need to calculate the second moment of interactions:

$$\langle a_{ij}^2 \rangle = \frac{1}{S(S-1)} \sum_{i \neq j} \sum_{\alpha} R_{i\alpha} R_{j\alpha} \sum_{\beta} R_{i\beta} R_{j\beta} = \frac{1}{S(S-1)} \sum_{\alpha\beta} \sum_{i} R_{i\alpha} R_{i\beta} \sum_{j} R_{j\alpha} R_{j\beta}$$
$$= \frac{1}{S(S-1)} \left( \sum_{\alpha \neq \beta} (\sum_i R_{i\alpha} R_{i\beta})^2 + \sum_{\alpha} (\sum_i R_{i\alpha}^2)^2 \right) = M(M-1)\mu_1(R)^4 + M\mu_2(R)^2 \tag{12}$$

where we consider contributions from independent and non-independent growth rates separately. Putting this together, we get the variance of interactions relating to the moments of growth rate distribution:

$$\kappa_2(a) = \sigma_a^2 = \langle a_{ij}^2 \rangle - \langle a_{ij} \rangle^2 = M(\mu_2(R)^2 - \mu_1(R)^4) \tag{13}$$

While we obtain the mean and variance, to define the skew, we need the third cumulant defined as $\kappa_3(a) = \langle a_{ij}^3 \rangle - 3\langle a_{ij}^2 \rangle \langle a_{ij} \rangle + 2\langle a_{ij} \rangle^3$. The calculations for this proceed along similar lines as those for variance. We highlight the important steps below for $\langle a_{ij}^3 \rangle$

$$\langle a_{ij}^3 \rangle = \frac{1}{S(S-1)} \sum_{i \neq j} \sum_{\alpha\beta\gamma} R_{i\alpha} R_{i\beta} R_{i\gamma} R_{j\alpha} R_{j\beta} R_{j\gamma}. \tag{14}$$

This can be split into three contributions depending on the values of $\{\alpha, \beta, \gamma\}$: $\alpha \neq \beta \neq \gamma$ having $M(M-1)(M-2)$ terms each equal to $\mu_1(R)^6$; combinations of $\alpha = \beta \neq \gamma$ with $3M(M-1)$ terms each contributing $\mu_2(R)^2\mu_1(R)^2$; and $\alpha = \beta = \gamma$ with $M$ terms of $\mu_3(R)^2$. Putting these terms together, we see that the third interaction cumulant is given by

$$\kappa_3(a) = M \left( \mu_3(R)^2 - 3\mu_2(R)^2 \mu_1(R)^2 + 2\mu_1(R)^6 \right) \tag{15}$$

allowing us to calculate the skew given by the ratio of cumulants,

$$\text{skew } s \equiv \frac{\kappa_3(a)}{\kappa_2(a)^{3/2}}. \tag{16}$$

Introducing cross-feeding complicates the model: we introduce dependencies between different resources in the steady state based on the stoichiometric matrix leading to the quasi-stationary resource concentrations depending on all resources. However, few simplifications are still possible. Consider the model where, for simplicity, we assume that all species leak at the same fraction $l$:

$$\dot{n}_i = n_i \left( \sum_{\alpha} (1-l) R_{i\alpha} c_\alpha - \delta \right) \tag{17}$$
$$\dot{c}_\alpha = c_\alpha (s - c_\alpha) - \sum_i R_{i\alpha} c_\alpha n_i + l \sum_{i\beta} n_i c_\beta D_{\alpha\beta} R_{i\beta} \tag{18}$$

At quasi-stationarity, i.e., $\dot{c}_\alpha = 0 \; \forall \alpha$,

$$0 = \sum_\beta c_\beta \left[ \delta_{\alpha\beta} \left( s - c_\beta - \sum_i n_i R_{i\beta} \right) + (1 - \delta_{\alpha\beta}) \left( \sum_i l n_i R_{i\beta} D_{\alpha\beta} \right) \right] \tag{19}$$

where $\delta_{\alpha\beta}$ is the Kronecker delta, which is one when $\alpha = \beta$ and zero otherwise. Note that this equation is of second order and hence an easy closed form solution is not feasible. Instead, we consider low leakage rates, which then allows us to perturbatively write the solution as:

$$c_\alpha(l) = c_{0\alpha} + l c_{1\alpha} + \mathcal{O}(l^2). \tag{20}$$

We know that $c_{0\alpha}$ corresponds to the quasi-stationary solution in the purely competitive case where $l = 0$. Then, we substitute the perturbative equation back in the solution and match the terms at $\mathcal{O}(l)$ to obtain the following:

$$c_{1\alpha} \left( s - 2c_{0\alpha} - \sum_i n_i R_{i\alpha} \right) = - \sum_{i\beta} n_i R_{i\beta} D_{\alpha\beta} c_{0\beta}. \tag{21}$$

By recognizing $c_{0\alpha} = s - \sum_i n_i R_{i\alpha}$, we obtain the first order correction

$$c_{1\alpha} = \sum_{i\beta} n_i R_{i\beta} D_{\alpha\beta} \frac{c_{0\beta}}{c_{0\alpha}}. \tag{22}$$

Note that in doing so, we have assumed that all resources are supplied even though there is leakage. If not, then $c_{0\alpha} = 0$ for unsupplied resources. This correction can be taken into account by separately calculating the perturbative expansion for supplied and non-supplied resources, but our following assumption would result in a similar final equation. We assume that the fraction $\frac{c_{0\beta}}{c_{0\alpha}}$ is of the same order and then equal on average. This corresponds to taking their mean-field limit and we ignore the context-dependence arising even in a low-leakage perturbative solution. Therefore, our quasi-stationary resource concentration for low-leakage values is given by

$$c^*_\alpha \approx s - \sum_i n_i \left( R_{i\alpha} - l \sum_\beta R_{i\beta} D_{\alpha\beta} \right). \tag{23}$$

This also gives us the correction to interactions caused by leakage

$$a_{ij}(l) \approx (1-l) \sum_\alpha R_{i\alpha} R_{j\alpha} - (1-l) l \sum_{\alpha\beta} R_{i\alpha} D_{\alpha\beta} R_{j\beta} \tag{24}$$

which immediately points to the fact that we cannot get arbitrary large and positive interaction values since it is always in a tradeoff with consumption, leading to limits on positive interaction. Now, as we did with the quasi-stationary resource concentration, we consider interaction terms upto $\mathcal{O}(l)$, and define two matrices for ease of calculations:

$$a_{ij}(l) \approx A_{ij} - l(A_{ij} + B_{ij}) \tag{25}$$

where $A_{ij} = \sum_\alpha R_{i\alpha} R_{j\alpha}$ is the competitive interaction and $B_{ij} = \sum_{\alpha,\beta} R_{i\alpha} D_{\alpha\beta} R_{j\beta}$ is the correction coming from cross-feeding. We proceed in a manner similar to before: compute the different powers of $a_{ij}$, take the averages to get the moments, and then calculate the cumulants till first order in $l$. Upon calculating the different possibilities of what indices are equal to what when multiple summations are present and simplifying the expressions, we obtain these approximations for the perturbative expansion of interaction cumulants in the presence of cross-feeding:

$$\kappa_1(l) = \kappa_1(A) - l\left[\kappa_1(A) + \kappa_1(B)\right] + \mathcal{O}(l^2) \tag{26}$$

$$\kappa_2(l) = \kappa_2(A) - 2l\left[\kappa_2(A) + Cov(A,B)\right] + \mathcal{O}(l^2) \tag{27}$$

$$\kappa_3(l) = \kappa_3(A) - 3l\left[\kappa_3(A) + Cov(AB,A) - \kappa_1(B)\kappa_2(A) - \kappa_1(A)Cov(A,B)\right] + \mathcal{O}(l^2) \tag{28}$$

with explicit expression of the different moments involved being

$$\begin{aligned}
\mu(A) =& M\mu(R)^2 \\
\mu(A^2) =& M(M-1)\mu(R)^4 + M\mu(R^2)^2 \\
\mu(A^3) =& M(M-1)(M-2)\mu(R)^6 + 3M(M-1)\mu(R^2)^2\mu(R)^2 + M\mu(R^3)^2 \\
\mu(B) =& M(M-1)\mu(R)^2\mu(D) \\
\mu(B^2) =& M(M-1)\mu(R^2)^2\mu(D^2) + 2M(M-1)(M-2)\mu(R^2)\mu(R)^2\mu(D)^2 \\
&+ M(M-1)(M-2)(M-3)\mu(R)^4\mu(D)^2 \\
\mu(AB) =& M(M-1)(M-3)\mu(R)^4\mu(D) + 2M(M-1)\mu(R^2)\mu(R)^2\mu(D) \\
\mu(A^2B) =& 2M(M-1)\mu(R^3)\mu(R^2)\mu(R)\mu(D) + 2M(M-1)\mu(R^2)^2\mu(R)^2\mu(D) \\
&+ 4M(M-1)(M-2)\mu(R^2)\mu(R)^4\mu(D) + M(M-1)(M-2)(M-3)\mu(R)^6\mu(D)
\end{aligned}$$

While we do obtain an expression for skew this way by including cross-feeding, the calculation is not strictly correct since this does not take into account the strong context dependence that can arise at higher leakage values. Furthermore, if we switch from biotic supply of resources to abiotic supply, then, the quasi-stationary concentration of the resources cannot be so easily determined, and even the purely competitive communities will have interactions that are context-dependent. Despite these assumptions and approximations, the obtained skew qualitatively mimics the simulation observations in Fig

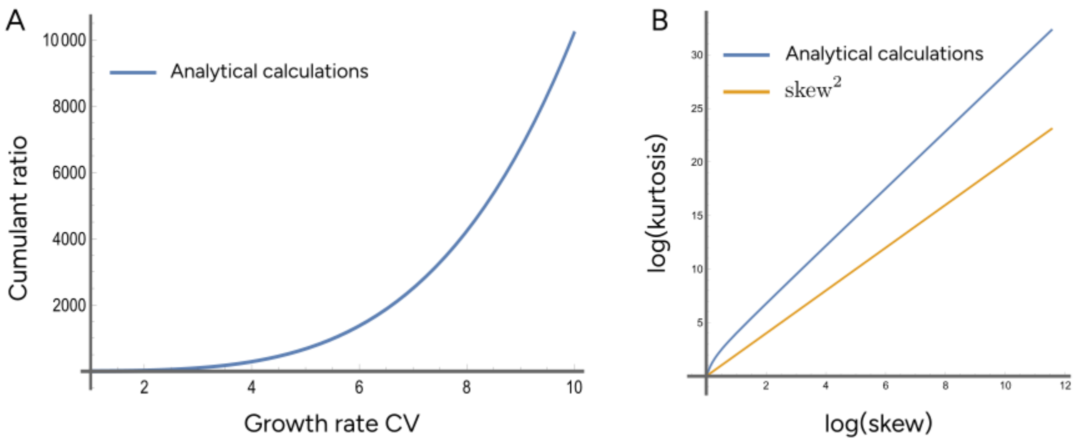


FIG. 4. Skewed and heavy tailed interaction distributions. **A)** Cumulant ratio ($\kappa_1\kappa_3/\kappa_2^2$) against growth rate coefficient of variation (CV) in the purely competitive case showing a consistent increase. A gamma distribution would have this value constant ($= 2$). **B)** Plot of log of kurtosis against log of skew in the competitive case showing that kurtosis values don't scale with skew as one would expect from a gamma distribution (kurtosis $\propto$ skew$^2$), indicating more presence of strong outliers, i.e., a heavy tailed distribution. For both plots, $M = 100$ and growth rates are from a lognormal distribution with mean $1/M$ and standard deviation set by the CV.

1D of the main text - that increasing growth rate heterogeneity and leakage fractions increase the skew. These calculations and the associated plots have been provided in the Mathematica notebook `skew_calculations.nb`

Upon seeing that skewness increases both with leakage and with competition, we can ask two questions: do the cumulants of the interaction show properties similar to any known distribution? how strong are the outliers? For the first, we look at the cumulant ratio: $\kappa_1\kappa_3/\kappa_2^2$. This is basically the ratio of skewness to the coefficient of variation of the interactions. We compute this for the competitive case (by setting $l = 0$) and plot it against the growth rate heterogeneity (CV). Figure 4A shows that this cumulant ratio increases with growth rate CV. For a gamma distribution, this ratio is exactly equal to 2. This difference from a constant value is consistent with distributions such as lognormal, which have stronger outliers than the gamma. This can be further investigated. Upon calculating the kurtosis for the competitive case and plotting it against skew, we see in Fig 4B that on a log-log scale, the kurtosis grows faster than $skew^2$. The scaling of $skew^2$ is again expected from a gamma distribution and a faster scaling indicates that for the same skew value, there is higher kurtosis, i.e., there are more outlier values. These plots indicate that interaction distribution from consumer-resource model is likely a heavy-tailed one, of which lognormal distribution is an example.

#### 6. INTERACTION MATRICES FROM DATA

For the synthetic duckweed community [5], interaction matrices (shown in Fig 5A) estimated from the Maynard et al procedure [6] produce a good estimate of the abundance of drop-in communities ($R^2 = 0.77$), shown in Fig 5B. We do not consider the correlated lognormal here since with only seven species, and hence 42 interspecific interactions, the inferred value might not be a good estimator of actual correlation (estimated correlation is -0.11 with a standard error of $\pm$ 0.15).

For the *in vitro* gut community, we show in Figure 5C the preferences of the effective resource groups inferred from spent media metabolomics as calculated in Ho et al [7]. We see that species 3, 6, and 14 (corresponding to *Bt*, *Pd* and *Csc* in the data, see Methods) all have a unique resource group that is only consumed by that species and no other. This means that according to the inferred growth rate matrix, these species do not interact with any others because no niche is shared with the rest of the community. If we were to include them in the EO framework to obtain an effective interaction matrix, they would only have zeros in their interspecific interaction values. Hence, we choose to drop these species to estimate the interactions. In doing so, we are likely to underestimate how weak interactions are on average. Ideally, the interactions estimated after dropping these species would be used to predict diversity of the 11 drop-one-out communities of the 12-member subcommunity that do not contain the three species (*Bt, Pd,* and *Csc*). However, the experiments do not include such cases and we only have the drop-one-outs from the full 15 member community. Hence, we attempt to predict the same, while acknowledging that our informed interactions are likely stronger than in reality.

To estimate the interactions, we run a chemostat model with a dilution rate of 0.11, which is equal to $-\log(D)/\tau$ where $D$ is the experimental dilution factor, i.e. 1:200 and $\tau = 48h$ is the time between dilutions. For the supply of resources, Ho et al [7] also estimate the initial serial dilution concentrations of their effective resource groups. Hence,

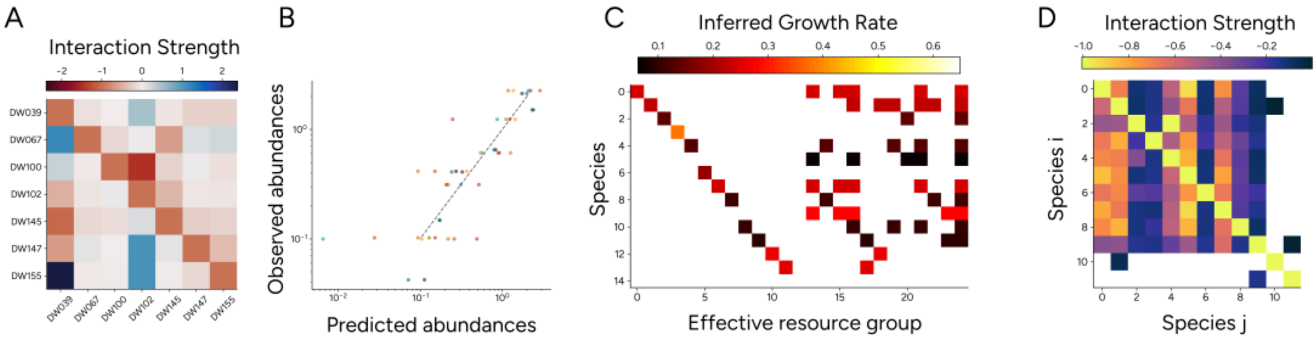


FIG. 5. Interaction matrices obtained from data **A)** Duckweed synthetic community interactions inferred from subcommunity abundances [5]. **B)** using the interaction matrix in panel A to predict out-of-sample abundances produces good correlation between predictions and experimental observations validating the obtained interaction matrix ($R^2 = 0.77$). **C)** Effective resource groups of *in vitro* human gut community inferred from metabolomics of spent media as reported in [7] with heatmap showing the growth rates on the specific groups. Three species (numbered 3, 6, and 14) don't share any resource group with the other species making them completely non-interacting with the rest of the community. **D)** Inferred interaction matrix using EO framework with the growth rate matrix in panel C, ignoring the non-interacting species still produces sparse values due to two species sharing only one other resource group with the community. This sparsity is included when making predictions in the main text (see Methods).

this competitive chemostat model has all relevant parameters directly obtained from experimental protocols or previously reported values. When we consider the interaction matrix after dropping the inferred non-interacting species, we still observe many zeros caused by two species (Fig 5D). These species have less resource overlap, but because they are not completely non-interacting from the rest of the community, we instead choose to consider them but take the sparsity into account while generating the informed gLV models (see Methods).

---

[1] Richard R Stein, Vanni Bucci, Nora C Toussaint, Charlie G Buffie, Gunnar Rätsch, Eric G Pamer, Chris Sander, and Joao B Xavier. Ecological modeling from time-series inference: insight into dynamics and stability of intestinal microbiota. *PLoS computational biology*, 9(12):e1003388, 2013.
[2] Anna S Weiss, Anna G Burrichter, Abilash Chakravarthy Durai Raj, Alexandra von Strempel, Chen Meng, Karin Kleigrewe, Philipp C Münch, Luis Rössler, Claudia Huber, Wolfgang Eisenreich, et al. In vitro interaction network of a synthetic gut bacterial community. *The ISME journal*, 16(4):1095–1109, 2022.
[3] Jared Kehe, Anthony Ortiz, Anthony Kulesa, Jeff Gore, Paul C Blainey, and Jonathan Friedman. Positive interactions are common among culturable bacteria. *Science advances*, 7(45):eabi7159, 2021.
[4] Martin Schäfer, Alan R Pacheco, Rahel Künzler, Miriam Bortfeld-Miller, Christopher M Field, Evangelia Vayena, Vassily Hatzimanikatis, and Julia A Vorholt. Metabolic interaction models recapitulate leaf microbiota ecology. *Science*, 381(6653):eadf5121, 2023.
[5] Hidehiro Ishizawa, Yosuke Tashiro, Daisuke Inoue, Michihiko Ike, and Hiroyuki Futamata. Learning beyond-pairwise interactions enables the bottom–up prediction of microbial community structure. *Proceedings of the National Academy of Sciences*, 121(7):e2312396121, 2024.
[6] Daniel S Maynard, Zachary R Miller, and Stefano Allesina. Predicting coexistence in experimental ecological communities. *Nature ecology & evolution*, 4(1):91–100, 2020.
[7] Po-Yi Ho, Taylor H Nguyen, Juan M Sanchez, Brian C DeFelice, and Kerwyn Casey Huang. Resource competition predicts assembly of gut bacterial communities in vitro. *Nature Microbiology*, 9(4):1036–1048, 2024.
[8] Ryan L Clark, Bryce M Connors, David M Stevenson, Susan E Hromada, Joshua J Hamilton, Daniel Amador-Noguez, and Ophelia S Venturelli. Design of synthetic human gut microbiome assembly and butyrate production. *Nature communications*, 12(1):3254, 2021.
[9] Flor I Arias-Sánchez, Björn Vessman, Alice Haym, Géraldine Alberti, and Sara Mitri. Artificial selection improves pollutant degradation by bacterial communities. *Nature communications*, 15(1):7836, 2024.
[10] Ewa Merz, Riley J Hale, Erik Saberski, Kasia M Kenitz, Melissa L Carter, Jeff S Bowman, and Andrew D Barton. Temperature alters interactions and keystone taxa in the marine microbiome. *The ISME Journal*, 20(1):wraf287, 2026.
[11] Oliver J Meacock and Sara Mitri. Environment-organism feedbacks drive changes in ecological interactions. *Ecology letters*, 28(1):e70027, 2025.
[12] James P O'Dwyer. Whence lotka-volterra? conservation laws and integrable systems in ecology. *Theoretical Ecology*, 11(4):441–

452, 2018.

[13] Robert MacArthur. Species packing and competitive equilibrium for many species. *Theoretical population biology*, 1(1):1–11, 1970.

[14] Thomas Koffel, Tanguy Daufresne, and Christopher A. Klausmeier. From competition to facilitation and mutualism: a general theory of the niche. *Ecological Monographs*, 91(3):e01458, 2021.

[15] Yizhou Liu, Jiliang Hu, Hyunseok Lee, and Jeff Gore. Complex ecosystems lose stability when resource consumption is out of niche. *Physical Review X*, 15(1):011003, 2025.

[16] Akshit Goyal, Jason W. Rocks, and Pankaj Mehta. Universal Niche Geometry Governs the Response of Ecosystems to Environmental Perturbations. *PRX Life*, 3(1):013010, February 2025.

[17] Michael Mustri, Quqiming Duan, and Samraat Pawar. Accuracy of the Lotka-Volterra Model fails in strongly coupled microbial consumer-resource systems, February 2025.

[18] Lloyd N Trefethen and David Bau. Numerical linear algebra. chapter 12. SIAM, 2022.
